# A minimally guided organoid model for cross-species comparisons of cerebellar development

**DOI:** 10.1101/2024.10.02.616236

**Authors:** Luca Guglielmi, Daniel Lloyd-Davies-Sánchez, José González Martínez, Madeline A. Lancaster

## Abstract

The human cerebellum has undergone significant evolutionary expansion compared to other species, contributing to both motor and cognitive skills. However, the mechanisms underlying this process remain largely unknown as interrogating human cerebellar development alongside other species has to date been unfeasible. To address this, we developed a minimally guided cerebellar organoid protocol that combines unguided neural induction with precise temporal calibration of posteriorizing morphogens. This approach effectively overrides default telencephalic fate in cerebral organoids and induces stable cerebellar identities. Cerebellar organoids derived from both human and mouse ESCs exhibit species-specific size differences at comparable developmental stages and show robust induction of cerebellar master regulators and progenitor cell types. This model provides a powerful tool for investigating the mechanisms underlying cerebellar development in the context of both evolution and disease.

## Introduction

The cerebellum contains 80–90% of the neurons in the adult brain and plays a crucial role in both motor control and cognitive functions^1^. Cerebellar development is initiated in the most anterior region of the hindbrain, rhombomere 1 (r1), adjacent to the midbrain-hindbrain boundary (MHB) organizer, which separates the anterior from the posterior brain^2,3^. The induction of the MHB is established through the antagonistic interaction between *OTX2* and *GBX2*, which define the anterior and posterior epiblast, respectively^4^. Once the MHB is formed, ligands secreted from the organizer, such as WNTs and FGFs, promote cerebellar competency^5,6^ (Figure 1A). These signals, together with master regulators such as *PAX2, EN1*, and *EN2* ^5,7–11^, initiate a morphogenetic cascade guiding the formation of cerebellar progenitors and mature neurons. While these early patterning events are broadly conserved across vertebrates, the cerebellum underwent rapid expansion during human evolution, significantly contributing to the overall increase in brain size^12,13^. In primates, cerebellar expansion occurred in tandem with neocortical growth^14,15^. However, humans and other apes deviate from this pattern, exhibiting a disproportionately large cerebellum relative to neocortex size^13,16^. This is in line with the key role of the cerebellum in modulating technical abilities, communication and emotional processing^13,16–18^. However, investigating mechanisms underlying these human-specific features has been hampered by the lack of tractable cerebellar models for humans and other species.

**Figure 1.**
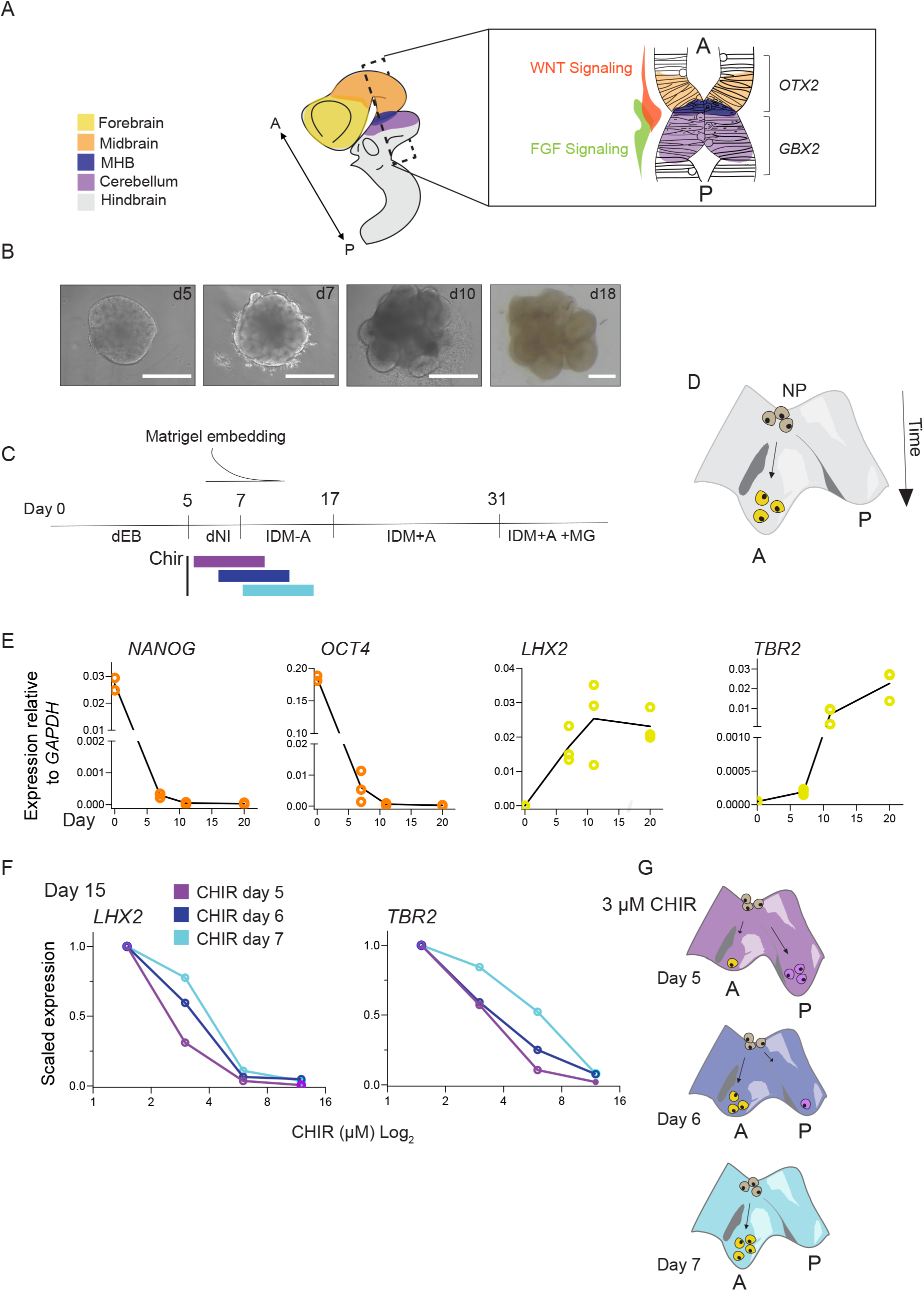
Telencephalic identities are suppressed by early WNT signaling activation. **A)** Schematics representing A/P patterning of the human embryo brain, where different A/P regions are color coded. Cross section focuses on signaling at the MHB and induction of cerebellar master regulators at the posterior end of the boundary. A, anterior; P, posterior. **B)** Brightfield images representing unguided organoids at the indicated stages. Scalebars 400μm. **C)** Unguided protocol timeline using chemically defined embryo body (dEB) media and neural induction media (dNI). IDM-A, improved differentiation media - viatamin A; IDM+A, improved differentiation media + vitamin A. **D)** Illustration depicting neural progenitor differentiation under unguided conditions; uncommitted progenitor most likely default towards telencephalic fates instead of posterior identities. NP, neuronal progenitors. **E)** qPCR for pluripotency markers and anterior brain markers over time. Each circle represents an independent batch and experiment. For all genes, qPCR data are the result of two independent experiments for day 0 and three independent experiments for day 11 and 20. **F)** qPCR for *LHX2* and *TBR2* at day 15 upon CHIR dose response using either 1.5, 3, 6 and 12 μM. Day of treatment is indicated by different colours. qPCR data are the result of one independent experiment and relative expression values across different days and doses were normalized on the 1.5 μM dose for each gene. **G)** As for (E), illustration represent neural progenitor differentiation; unlike for unguided conditions, anterior fate induction is reduced when 3μM CHIR is used at day 5.

Alongside these evolutionary aspects, cerebellar growth is disrupted in several human neurodevelopmental conditions, including Dandy-Walker malformation^19^, Joubert syndrome^20^, and ataxia telangiectasia^21^, as well as in medulloblastoma^22,23^ the most common paediatric brain cancer^24^. While animal models have been instrumental in advancing our understanding of these diseases, they often do not fully replicate human-specific pathophysiological features. Thus, comparing the effects of disease determinants between humans and other species is essential to uncover the mechanisms underlying human-specific penetrance or expressivity of disease.

To date, a few cerebellar organoid models have been developed^25–27^which have opened up the possibility to explore development of different cerebellar neuron types. These models rely heavily on guided differentiation protocols, allowing for robust determination of the different cerebellar cell types. An alternative, minimally guided method with minimal to no extrinsic manipulation has not yet been described but would enable investigation of species-specific differences in intrinsic developmental programs that may underlie differences in tempo and/ or the topology of developmental transitions.

Notably, unguided cerebral organoids have provided crucial insights into the mechanisms underlying evolutionary differences in brain size ^28–30^ as well as into human diseases ^31,32^. In these models, neural progenitors default towards telencephalic fates and self-organize into distinct cortical progenitor layers^33^. While these models well recapitulate anterior brain development, spontaneous acquisition of posterior identities like the cerebellum occurs at a negligible frequency.

To address this issue, we combine unguided neural induction with temporally calibrated WNT and FGF signalling activation to override telencephalic fates and stabilize cerebellar identities. This approach, which relies on minimal and conserved cerebellar determinants, enables cross-species applicability. Indeed, cerebellar organoids derived from both human and mouse recapitulate species-specific differences in size at comparable developmental stages, they also display robust induction of cerebellar master regulators and the generation of cerebellar progenitor cell types with timing that closely mirrors in vivo development. This platform provides a powerful tool for cross-species comparisons and investigation of human-specific features in cerebellar development across evolution and disease.

## Results

### Overriding the default telencephalic program to induce posterior brain identities

Under unguided conditions, hESCs exit pluripotency and differentiate into neuroepithelial progenitors as the default, predominantly acquiring telencephalic identities^29,34^. We hypothesized that providing caudalizing signals after neuroepithelial identity is determined but before telencephalic fate becomes irreversible could promote posterior identities. To test this hypothesis, we first investigated the timing of hESC exit from pluripotency and the initiation of forebrain marker expression. Using a defined media composition, we generated unguided brain organoids from H9 hESCs (Figure 1B-D). We then examined the expression of pluripotency markers *NANOG* and *OCT4*, alongside early forebrain markers *LHX2* and *TBR2*, a marker of cortical intermediate progenitors, during the first 20 days of differentiation (Figure 1E). Notably, *NANOG* and *OCT4* expression were significantly reduced by day 7 and absent by day 11 (Figure 1E). In contrast, *LHX2* and *TBR2* expression were already detectable at day 7 and increased progressively (Figure 1E), suggesting that by day 7, the forebrain specification program was already underway. Based on these findings, we aimed to introduce caudalizing cues before this critical time point. Given the conserved role of WNT signaling in early anterior-posterior (A/P) patterning, where early pathway activation prevents forebrain development^35^ and inhibition suppresses formation of midbrain and hindbrain^36,37^; we investigated whether activation of WNT signaling just prior to day 7 would prevent telencephalic identities while still allowing the earlier neuroepithelial establishment. To determine the correct time window and dosage, we performed a dose response experiment using the WNT signalling activator CHIR at day 5, 6 and 7 (Figure 1C and F). CHIR was maintained for 4 days and expression of *LHX2* and *TBR2* was assessed at day 15 using qPCR (Figure 1F).

Upon treatment, expression of both *LHX2* and *TBR2* scaled with the CHIR dosage, with no detectable expression for either marker at the 12 μM dose, regardless of treatment onset (Figure 1F). However, at this high concentration, organoid morphology was compromised (Figure S1). We therefore sought to identify a dosage that would both maintain proper neuroepithelial morphology and effectively suppress forebrain identities ^36^. Notably, CHIR administration at lower doses revealed that treatment initiated on day 5 was more effective at suppressing anterior marker genes compared to treatments started on days 6 or 7 (Figure 1F). For instance, the 3μM dose had little effect on *LHX2* and *TBR2* expression when given at day 6 or 7 but reduced expression of both markers about twofold when given at day 5 without gross morphological abnormalities (Figure 1F and S1). This further supports the notion that, by day 7, neural progenitors are already committed to anterior identities and this specification can be prevented by early posteriorizing cues (Figure 1G). With this temporal and dose information, we next sought to steer progenitor fates towards cerebellar identities.

### Combinatorial WNT and FGF signalling activation stabilizes cerebellar identities

In the embryo, cerebellar development relies on the coordinated activity of WNT and FGF signaling at the MHB (Figure 1A). WNT signaling is essential for the formation and maintenance of the MHB^36–39^ while FGF signaling plays a crucial inductive role in establishing cerebellar competency within neuroepithelial progenitors^5,8,40,41^. The synergy between these pathways is critical, as FGF signaling on its own can promote telencephalic identities during early brain development^42,43^. Although these relationships are well established in vivo, the situation in vitro remains less clear. Since the pioneering work of Yoshiki Sasai and collaborators ^44^ cerebellar identities have been successfully generated in both 2D and 3D systems. These protocols typically employ dual SMAD inhibition followed by either combined FGF and WNT activation ^27,45^ or FGF signalling alone^25,44,46,46^, showing that both approaches can effectively promote cerebellar identities.

To address this in our system and preserve as much as possible the intrinsic developmental program, we combined unguided neural induction (i.e., without dual SMAD inhibition) with treatment at day 5 using either 3 μM CHIR in combination with FGF2 or FGF2 alone (Figure 2A). Although FGF8 is the main inducer of cerebellar fates in vivo, FGF2 has been shown to be superior in vitro in inducing the expression of *EN2* and widely used to reproduce cerebellar identities ^25,44,45^.

**Figure 2.**
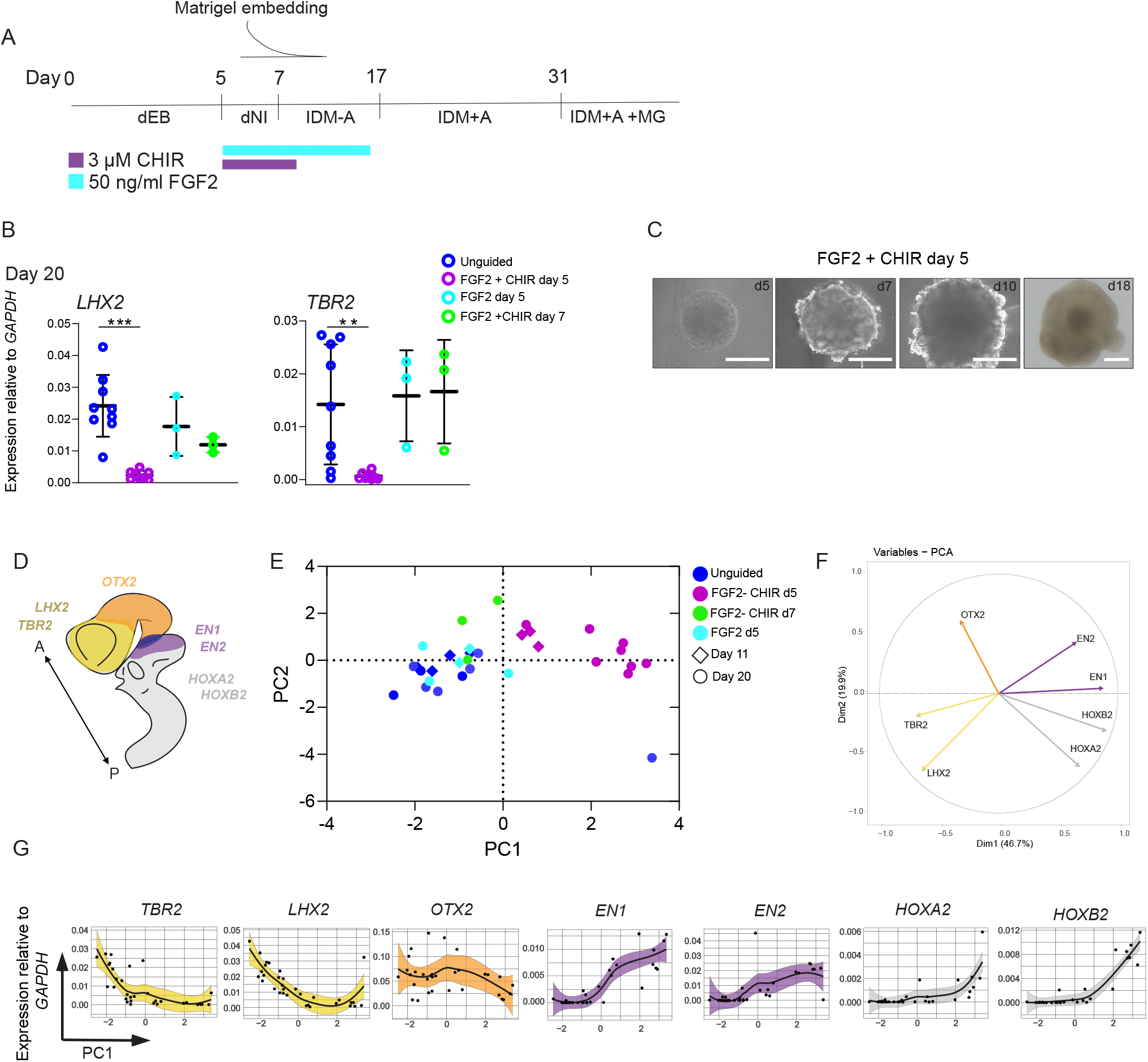
Caudalized organoids display expression of early cerebellar markers. **A)** Cerebellar protocol timeline displaying timing of CHIR and FGF2 treatment. dEB, defined embryo body media; dNI, defined neural induction media (dNI); IDM-A, improved differentiation media - viatamin A; IDM+A, improved differentiation media + vitamin A. **B)** qPCR for *TBR2* and *LHX2* at day 20. Each circle represents an independent batch and experiment for each condition. For the unguided condition, qPCR data are the result of nine independent batches and experiments, for FGF2+CHIR day 5, eight independent batches and experiments. Finally, for FGF2 Day5 and FGF2+CHIR day 7 data represent three independent batches and experiments. Means +/- SD are shown. Significance was tested between the unguided condition and the FGF2 + CHIR using a t test with Welch’s correction (*TBR2*, P= 0.0071; *LHX2* P= 0001). **C)** Brightfield images representing caudalized organoids at the indicated stages. Scalebars 400μm. **D)** Illustration representing the human embryo brain with anterior posterior regions alongside master regulator genes are color coded. **E)** PCA plot across different unguided and minimally guided conditions. Each dot represents an individual batch and experiment. PC1, Principal component 1; PC2, principal component 2. **F)** Variables plot showing contribution of the different gene variable in D to the axis of the PCA plot in E. **G)** Plots sowing relative expression values for the genes in D in respect to PC1, dots represent individual batches as in E and confidence interval is color coded based on D.

At first, we sought to explore whether addition of FGF2 alongside CHIR would enhance or reduce WNT mediated suppression of *LHX2* and *TBR2* observed at day 20. Notably expression of both genes was fully suppressed by combinatorial signalling activation, an effect that was much stronger compared to either CHIR or FGF induction alone (Figure 2B and Figure 1F). Interestingly, in line with previous results, expression of *LHX2* and *TBR2* was still present if combinatorial treatment was performed at day 7 instead of day 5 (Figure 2B). Notably, these minimally guided organoids displayed distinct morphological features compared to their unguided counterparts (Figure 1B and 2C), showing smaller buds and a few neural crest cells developing on their surface (Figure 2C); events which are developmentally coupled with posterior neural tube ^47^ and posterior neural plate border induction ^48^, respectively. Supporting this, the loss of anterior markers was accompanied by the induction of posterior brain markers *GBX2* and *PAX2* (Figure S2A and S2B). We also observed a mild reduction in the expression of the neural progenitor marker *SOX2* (Figure S2A), consistent with the presence of non-neural identities potentially coming from neural crest.

To gain further insight into the A/P positioning of minimally guided organoids we probed for expression of an array of key master regulator genes spanning from the anterior to the posterior end of the brain at days 11 and 20 (Figure 2D-E). Expression profiles across all batches were integrated using principal component analysis (PCA). Interestingly, organoids organized in PCA space with a distinct trajectory along PC1 (Figure 2E) which represented A/P axis. This was assessed by looking at PCA loadings (Figure 2F) as well as plotting relative expression values against PC1 (Figure 2G). Accordingly, unguided organoids predominantly clustered on the left side of the plot, driven by high expression of anterior markers *TBR2* and *LHX2*. Similarly, organoids treated only with FGF2 and CHIR+FGF2 at day 7 displayed a comparable pattern, with a few showing intermediate, midbrain-like identities (Figure 2E). In contrast, combinatorial treatment at day 5 had a stronger caudalizing effect resulting in a marked shift in organoid identities driven by robust induction of *EN1* and *EN2* and the anterior hindbrain markers *HOXA2* and *HOXB2* (Figure 2E). Thus, the combination of WNT and FGF activation can effectively override anterior identities and reproduce MHB signalling and expression of cerebellar master regulators. We next investigated whether this would also lead to the generation of cerebellar progenitors.

### Minimally guided cerebellar organoids exhibit cerebellar progenitor zones

The cerebellar primordium is subdivided into two distinct progenitor zones positioned along the dorsal-ventral axis: the ventricular zone (VZ) and the rhombic lip (RL). The VZ is located ventrally and is marked by *PTF1A* and *KIRREL2*. This region will give rise to Purkinje cells (PCs), deep cerebellar nuclei (DCN) and interneurons. Instead, the RL, which is more dorsal, is marked by *ATOH1* and gives rise to granule cells (GCs)^49–51^(Figure 3A). If this minimally guided approach effectively recapitulates early cerebellar induction, we should expect the emergence of these distinct progenitor zones over time in the organoids. To test this, we assessed expression of the distinct VZ progenitor markers *PTF1A* and *KIRREL2* as well as RL marker *ATOH1* in both unguided and minimally guided organoids at day 20 and day 35 (Figure 3B). In line with the above results, expression of these genes was uniquely induced upon early CHIR and FGF2 treatment and increased over time (Figure 3B). A similar trend was observed for induction of the post mitotic Purkinje cell progenitor marker *SKOR2* (Figure S2B).

**Figure 3.**
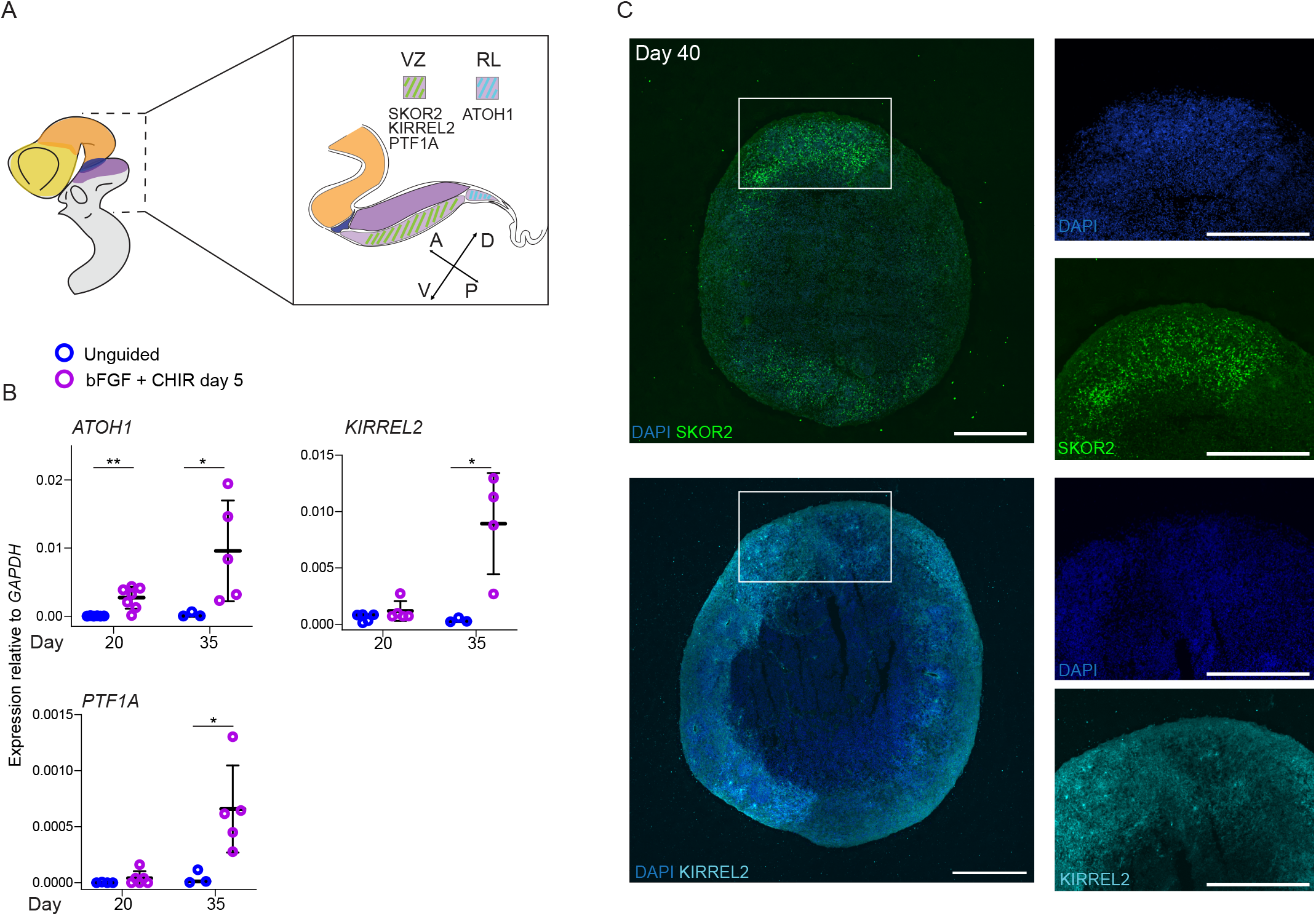
Minimally guided organoids express cerebellar progenitor markers. **A)** Schematics representing the human embryo brain. Cross section focuses on MHB and cerebellar primordium where different progenitor regions are color coded. VZ, ventricular zone; RL, rhombic lip. **B)** qPCR for *ATOH1, KIRREL2* and *PTF1A* on day 20 and 35. Each circle represents an independent batch and experiment for each condition. For *ATOH1, KIRREL2 and PTF1A* qPCR data are the result of at least three independent experiments for each condition and day. Means +/-SD are shown. Significance was tested using a t test with Welch’s correction (*ATOH1* day 20: P= 0.0044; day 35 P= 0.0472. *KIRREL2* day 20: P= 0.2250; day 35 P= 0.0408. *PTF1A* day 20: P= 0.1863; day 35: P= 0.0227). **C)** Immunostaining for SKOR2 and KIRREL2 on 12 μm human organoids slices at day 40. Scalebar 500 μm.

To complement these experiments, we performed immunostaining for the VZ markers KIRREL2 and SKOR2 at day 40, just after the onset of *SKOR2* expression (Figure S2B). In line with gene expression data, both markers were robustly expressed in minimally guided organoids (Figure 3C). Notably, *SKOR2* expression was organized in a radial pattern around the organoid, resembling radial migration of VZ progenitors in vivo ^52,53^(Figure 3C). Overall, these data show that alongside cerebellar competency, minimally guided organoids develop further into the cerebellar program by generating VZ and RL identities.

### Minimally guided induction of cerebellar identities translates across species

Given this protocol seems to recapitulate early cerebellar development in human organoids, we next explored whether this approach could also be applied to mouse ESCs and enable cross-species comparisons of cerebellar development. To achieve this, we made brain organoids using mouse E-14 ESCs. Notably, brain development in mouse occurs at a much faster pace compared to humans, therefore protocol steps were scaled according to mouse developmental tempo as previously performed ^54^. Similarly to our previous approach, we tested the effect of applying posteriorizing cues at different times during the organoid differentiation protocol. Application of CHIR + Fgf2 beginning at day 2 or 3 resulted in the visible development of mesenchymal cell identities (Figure S3A), suggesting a non-neuroepithelial identity. Instead, treatment from day 4 prevented this effect, leading to the formation of organoids which resembled the morphology observed in human (Figure 4A, B and Figure 2C).

**Figure 4.**
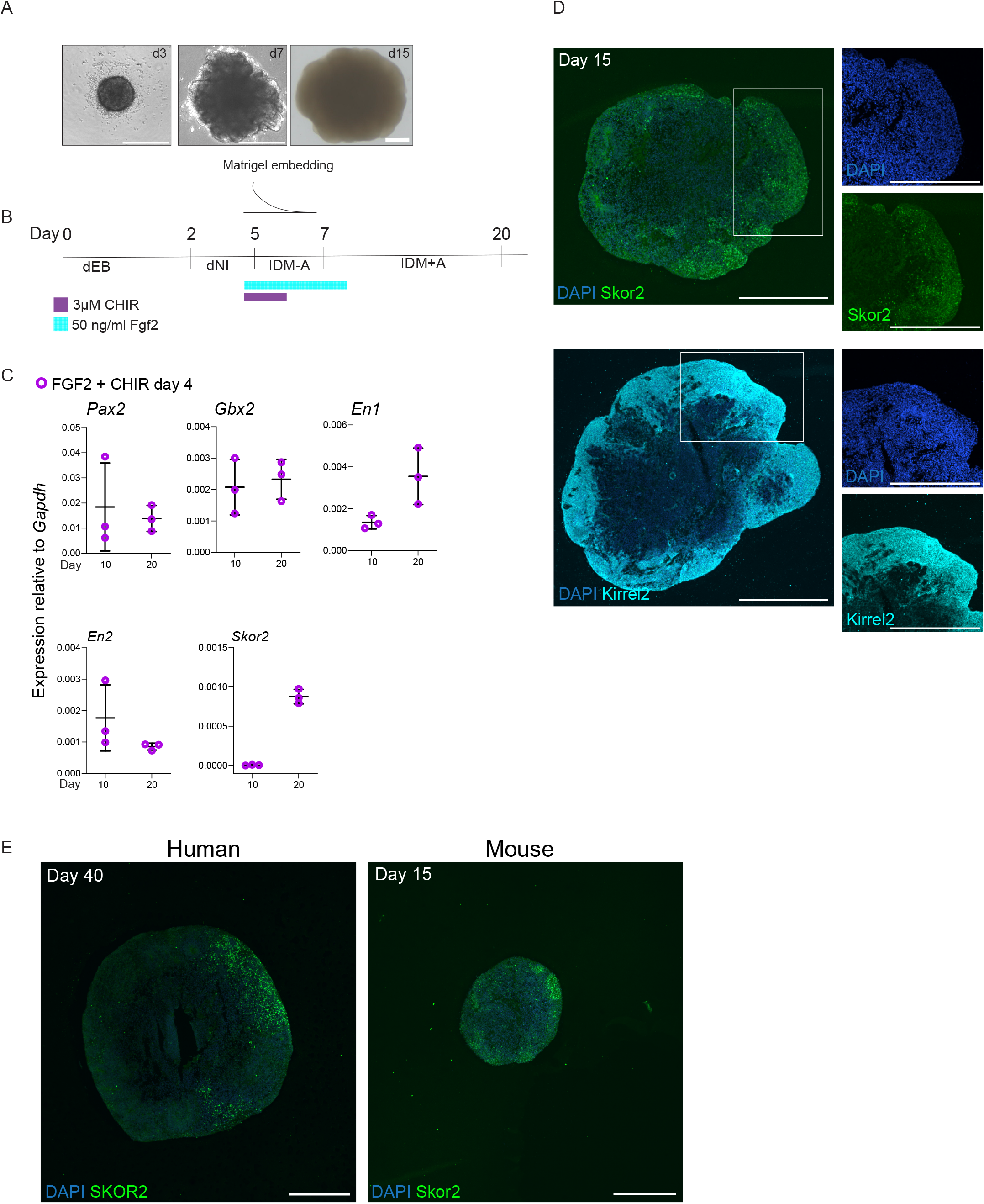
Inducing cerebellar identities in mouse organoids. **A)** Brightfield images representing caudalized mouse organoids at the indicated stages. Scalebars 400μm. **B)** Cerebellar protocol timeline adjusted for mouse developmental tempo timeline. dEB, defined embryo body media; dNI, defined neural induction media (dNI); IDM-A, improved differentiation media - viatamin A; IDM+A, improved differentiation media + vitamin A. **C)** qPCR for posterior markers in day 10 and 20 mouse organoids. Each circle represents an independent batch and qPCR data are the result of three independent experiments for each gene and each day. **D)** Immunostaining for Skor2 and Kirrel2 on 12 μm mouse organoids slices at day 15. Scalebar 500 μm. **E)** Immunostaining for SKOR2 and Skor2 for stage matched human and mouse organoids respectively. Scalebar 500 μm.

We next looked at expression of early cerebellar markers using qPCR at days 10 and 20. Intriguingly, as with human organoids, combinatorial Wnt and Fgf activation at day 4 suppressed the expression of anterior markers (Figure S3B) but induced robust expression of *Gbx2* alongside *Pax2, En1* and *En2* as well as the Purkinje progenitor marker *Skor2* (Figure 4C), without any detectable effect on *Sox2* expression (Figure S3B). However, we did not detect significant *Atoh1* expression (data not shown), suggesting that in mouse organoids, RL progenitors may be underrepresented. Notably, consistent with in vivo mouse development where *Skor2* expression begins around embryonic day 12 (E12) in the cerebellum ^44^, minimally guided mouse organoids did not express *Skor2* until after day 10 (Figure 4C). This shows that these organoids follow a developmental tempo comparable to that of in vivo embryogenesis.

To directly compare cerebellar progenitor marker expression between mouse and human organoids, we performed immunostaining for Skor2 and Kirrel2 at analogous developmental stages. Mouse organoids were stage matched based on the timing of *SKOR2* expression in human organoids (which begins around day 35, Figure S2B) and sliced at day 15. Immunostaining revealed a striking similarity with the human data in the expression pattern for both markers which displayed a comparable radial organization (Figure 4D). Intriguingly, in addition to these gene expression similarities, stage-matched mouse organoids were noticeably smaller than their human counterparts, resembling the in vivo differences in cerebellar size between the species (Figure 4E). This observation highlights how this minimally guided protocol recapitulates not only molecular features but also broad species-specific anatomical differences.

## Discussion

Overall, we developed a minimally guided cerebellar organoid model that effectively recapitulates early events in cerebellar development in both human and mouse ESCs. Building on unguided organoid models, we show that posterior brain identities can be induced using minimal signalling guidance. Indeed, by introducing posteriorizing cues before anterior commitment becomes irreversible, we successfully override anterior brain specification and promote cerebellar identities. This opens the possibility to explore specie-specific mechanisms underlying differences in cerebellar size. For instance, it has been shown that humans display a two-fold higher early Purkinje cell proportion compared to mouse, it would be intriguing to test whether this difference is dependent specie-specific delay in neurogenesis or due the unique presence of basal progenitors in the human cerebellum^51 50^.

Furthermore, the cross-species adaptability of this approach offers a valuable tool for investigating human-specific expressivity of cerebellar diseases. For instance, in ataxia-telangiectasia, the characteristic Purkinje cell degeneration observed in humans cannot be reproduced in mouse models^21^. Directly comparing how the same disease determinants affect both human and mouse organoids is key to understanding why human cells are vulnerable to degeneration while mouse cells are resistant. Uncovering these differences is crucial not only for identifying disease mechanisms but also for designing targeted therapeutic approaches.

## Methods

### Cell lines

Human ESCs (H9, female) and E-14 ESCs were used in this study. H9 (WA09) was purchased from WiCell, while E-14 were purchased from ATCC. Human and mouse ESCs used in this project were approved for use in this project by the UK Stem Cell Bank Steering Committee and approved by an ERC ethics committee and are registered on the Human Pluripotent Stem Cell Registry (hpscreg.eu). Cells were maintained in StemFlex for H9 (Thermo Fisher, A3349401) and Stemflex + mLIF (Sigma, ESG1106) for E-14. Both lines were grown on Matrigel (Corning, 356234) coated plates at 37 °C with controlled 5% CO2. Cells were passaged every 3-4 days using 0.7 mM EDTA.

### RNA isolation and cDNA preparation

For RNA extraction H9 hESCs and pooled human and mouse organoids were lysed though pipetting in 350μl lysis buffer (Cat# 74104, QUIAGEN RNeasy Mini Kit). For the human dataset 6 organoids for each batch were used for day 7 and 11, 4 organoids at day 20 and 3 organoids at day 35. For the mouse datasets 4 organoids were homogenized for day 10 and 3 organoids for day 20. RNA extraction was performed according to manufacturer’s instruction. To ensure complete removal of genomic DNA, the RNA extraction protocol was complemented with a DNase digestion as for manufacturer’s instruction (Cat# 79254, QUIAGEN RNase-Free DNase Set). RNA extracts were then processed for cDNA synthesis. This was performed using the AffinityScript kit (Agilent) as previously described ^55^. Briefly, 500 ng of RNA were retrotranscribed using random primers and the resulting cDNA was diluted 1:10 in deionized water for qPCR.

### qPCR and PCA analysis

All qPCRs were performed with the PowerUp SYBR Green Master Mix (ThermoFisher Scientific) with 300 nM of each primer and 2 μl of diluted cDNA. Fluorescence acquisition was performed on a ViiA 7 Real-Time PCR System (Thermo Fisher Scientific). Mouse and human primers are listed in Supplementary table 1. Quantification for relative gene expression was performed using the comparative Ct method and target gene expression was normalized to *GAPDH* for both human and mouse (*Gapdh*). For PCA analysis, relative expression values for each batch were scaled and visualized using the prcomp function using the Factoextra package in R (https://rpkgs.datanovia.com/factoextra/index.html). To visualize individual batched in PCA space, PC1 and PC2 coordinates were extracted and used for plotting in Prism10. Contribution of the different gene variables to PC1 and 2 was displayed using the fviz_pca_var function from the same package. Finally, plotting of relative expression value for each gene against PC1 was performed using the ggplot2 package (https://ggplot2.tidyverse.org/). Statistical analysis on qPCR data was performing using Prism10 and a t test with with Welch’s correction, P values are reported into figure legends.

### Fixing and sectioning

Human organoids at day 40 and mouse organoids at day 15 were washed in Phosphate Buffer Saline (PBS) and incubated in 4% PFA in phosphate buffer (PB) O/V at 4 °C. After fixation organoids were washed in PBS and moved in 30% sucrose/0.02% Sodium Azide (Sigma, S2002) in PB at 4°C until the tissue sank. The tissue was then embedded in 7.5% gelatin/30% sucrose in PB and frozen in a cold isopentane bath (Sigma, 277258) at -50°C. Samples were kept at -80°C until sectioning. Blocks were sectioned using a cryostat (Leica, CM1950) at 12 μm and sections were collected on charged slides (ThermoFisher Superfrost Plus, J1800AMNZ) and stored at -20°C until further processing.

### Immunohistochemistry and confocal imaging

Cryosections were rinsed in PBS 3×5 min. Slides were incubated in blocking buffer (0.1% Triton-X 100 with 4% donkey serum in PBS) for 1h at RT. Primary antibodies were diluted in blocking buffer at the following dilutions: rabbit anti-SKOR2 1:200 (Sigma, HPA046206), rabbit anti-KIRREL2 1:200 (Proteintech, 10890-I-AP). Primary antibody incubation was performed overnight at 4°C. The next day slides were washed in PBS and incubated with a secondary Donkey anti-rabbit Alexa Fluor 488 at 1:250 (Life Technologies) and DAPI 1:1000 in blocking buffer for 2h at RT. After PBS washes sections were let dry at RT and mounted in Prolong™ Diamond (Thermo Fisher Scientific, P36961) and stored at 4°C. Mounted sections were imaged with a Nikon W1 Spinning Disk scope using an a 10X objective.

### Organoids generation and neuronal patterning

To generate human and mouse unguided brain organoids single cell suspensions were obtained through incubations in Accutase (Sigma-Aldrich, A6964) for 4min at 37 °C. Next for both human and mouse ESCs, 4000 cells were seeded in U-bottom ultralow attachment 96 well plates (Corning, CLS7007) to generate embryo bodies (EBs) in defined EB media made of 2ml B27-A (Thermo Fisher Scientific, 12587010), 1 ml GlutaMAX (Thermo Fisher Scientific, 35050038), 1 ml of MEM-NEAA (Sigma, M7145), 96 ml DMEM-F12 (Thermo Fisher Scientific, 11330032) and supplemented with 50μM ROCK inhibitor Y27632 (Millipore, SCM075). EB media was kept from day 0 to day 5 for human organoids and from day 0 to day 2 for mouse organoids. Next EBs were moved in neural induction medium, composed of 1ml N2 supplement (Thermo Fisher Scientific, 17502048), 1 ml GlutaMAX (Thermo Fisher Scientific, 35050038), 1 ml of MEM-NEAA (Sigma, M7145), 100 μl Heparin solution (1mg/ml in PBS) in 96.9 ml DMEM-F12 (Thermo Fisher Scientific, 11330032). While in neural induction media, organoids were embedded in Matrigel droplets (Corning, 356234), this was done at day 6 for the human protocol and at day 4 for mouse and kept for 8 and 4 days respectively prior Matrigel removal using a fine scalpel. Human organoids were next incubated in improved differentiation medium without vitamin A (IDM-A) at day 7 as previously described ^56^ while for mouse this was done at day 5. As a final step, human organoids were moved to an orbital incubator at day 14 and bathed in improved differentiation medium plus vitamin A (IDM+ A) from day 18 until day 31, from this day on organoids were fed with IDM+A with 1 ml dissolved Matrigel per 50 ml media ^56^. Diversely, mouse organoids were moved to IDM+A at day 7 and to an orbital incubator at day 9 until sample collection. For the CHIR dose response, unguided human BOs were treated with either 1.5, 3, 6 or 12 μM CHIR (Tocris, 4423) and treatment was started at either day 5, 6 or 7 and maintained for 4 consecutive days. For this experiment organoids were embedded liquid Matrigel (5% in dNI media) instead of droplets which was added at day 5 and diluted through media change at each following day. For MHB patterning human organoids were treated with 50ng/ml human FGF2 (Human FGF-basic, Peprotech, 100-18B-100) from day 5 to day 14 and with 3 μM CHIR from day 5 to day 9, given FGF2 has a very short half-life at 37°C media was changed every day during the treatment. For mouse, 3 μM CHIR treatment was performed from day 4 alongside 50ng/ml Fgf2 (Murine FGF-basic, Peprotech, 450-33-100). CHIR treatment was maintained for two days while Fgf2 was maintained for three additional days.

## Acknowledgments

The authors would like to thank members of the Lancaster lab for their helpful feedback and discussions. We also thank the Light Microscopy facility of the MRC Laboratory of Molecular Biology. This work was supported by the Medical Research Council (MC_UP_1201/9) and a Vallee Scholars Award from the Vallee Foundation.

## Author contributions

L.G. conceived and designed the study. LG conducted experiments and analysed data with the help of D.L.S and J.G.M. LG wrote the manuscript. M.A.L. designed the study and supervised the project.

## Notes

### Competing Interest Statement

The authors have declared no competing interest.

