## Supplementary Material for "A minimally guided organoid model for cross-species comparisons of cerebellar development"

### Inventory supplementary information

**Figure S1. Effect of CHIR dose response on organoid morphology**

Related to Figure 1

**Figure S2. Combinatorial signaling induces robust expression of posterior marker genes**

Related to Figure 2

**Figure S3. Assessing expression of cerebellar markers in caudalized mouse organoids**

Related to Figure 4

**Supplementary table 1**

### Figure legends

#### Figure S1. Effect of CHIR dose response on organoid morphology

Brightfield images representing organoid batches on day 15 treated with different doses of the WNT activator CHIR on day 5, doses are indicated on the left-hand side.

#### Figure S2. Combinatorial signaling induces robust expression of posterior marker genes

**A)** qPCR for the gene set used for the PCA plot in figure 2E plus *GBX2* and *SOX2*. Each circle represents an independent batch and experiment for each condition. For the unguided condition, qPCR data are the result of nine independent batches and experiments for all genes beside *GBX2* and *SOX2* which are represented by six and eight experiments respectively. For the FGF2+CHIR day 5 condition, eight independent batches and experiments were performed for all genes beside six experiments for *GBX2* and seven experiments for *SOX2*. Means +/- SD are shown. Significance was tested using a t test with Welch's correction (*GBX2*, P= 0.035; *EN1*, P= 0.043; *EN2*, P< 0.0001; *HOXA2,* P< 0.56; *HOXB2*, P= 0.0063; *SOX2*, P= 0.0268. **B)** qPCR for *PAX2* at day 35 and *SKOR2* at day 11, 20 and 35.

**Figure S3. Assessing expression of cerebellar markers in caudalized mouse organoids**

**A)** Brightfield images showing mouse organoids treated with CHIR+ Fgf2 at day 2 and 3. Scalebar 500 µm**. B)** qPCR for *Sox2* and *Lhx2* on day 10. The data are the results of three independent batches and experiments for the unguided condition and four independent experiments for Fgf2 + CHIR at day 4.

Figure S1

Day 5 treatment

1.5 μM CHIR

3 μM CHIR

6 μM CHIR

12 μM CHIR

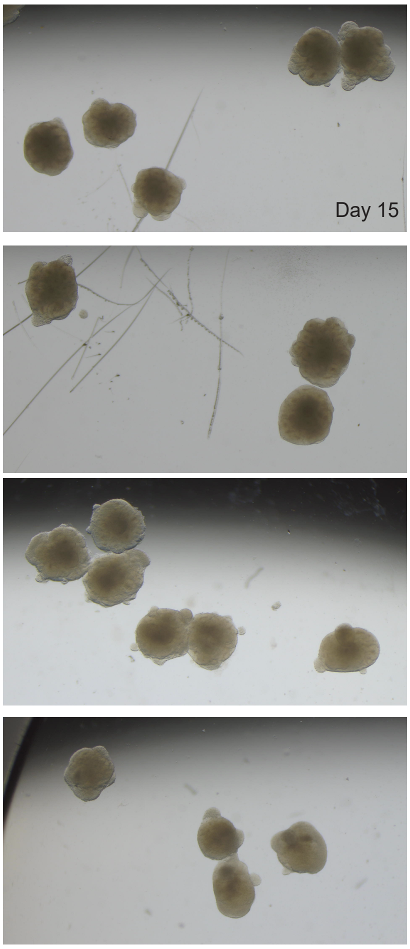

Figure S2

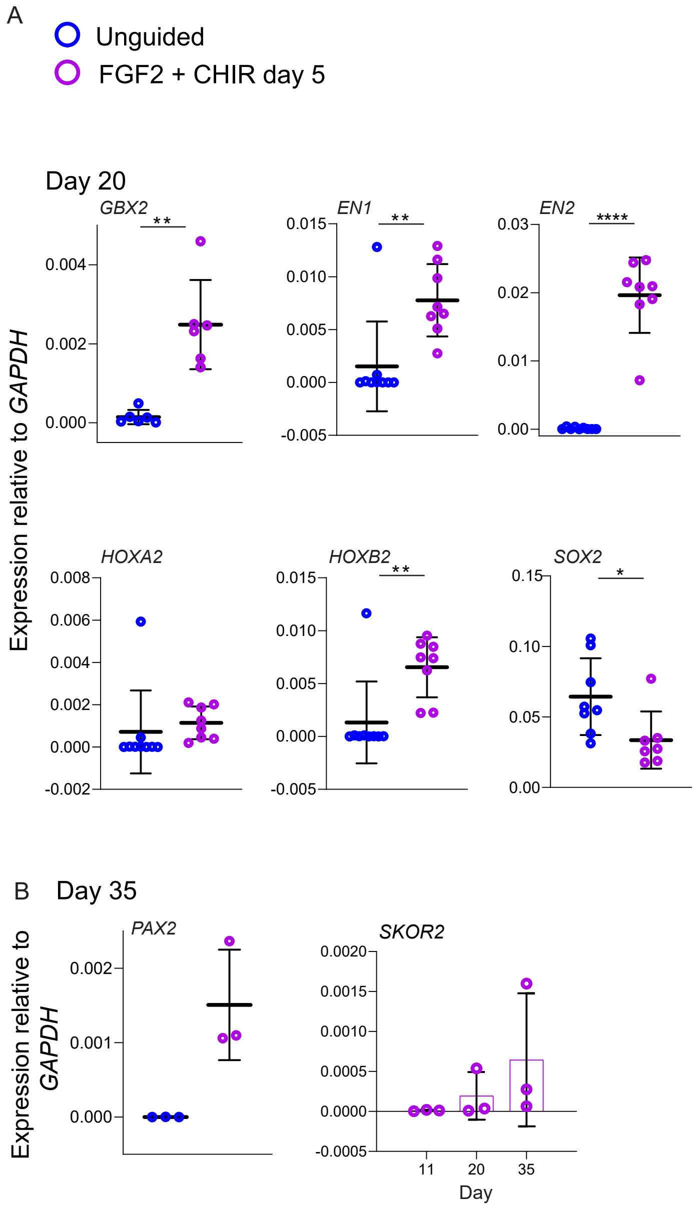
A
Unguided
FGF2 + CHIR day 5
Day 20
Expression relative to GAPDH
GBX2
**
0.004
0.002
0.000
EN1
**
0.015
0.010
0.005
0.000
-0.005
EN2
****
0.03
0.02
0.01
0.00
HOXA2
0.008
0.006
0.004
0.002
0.000
-0.002
HOXB2
**
0.015
0.010
0.005
0.000
-0.005
SOX2
*
0.15
0.10
0.05
0.00
B
Day 35
Expression relative to GAPDH
PAX2
0.002
0.001
0.000
SKOR2
0.0020
0.0015
0.0010
0.0005
0.0000
-0.0005
11
20
35
Day

Figure S3

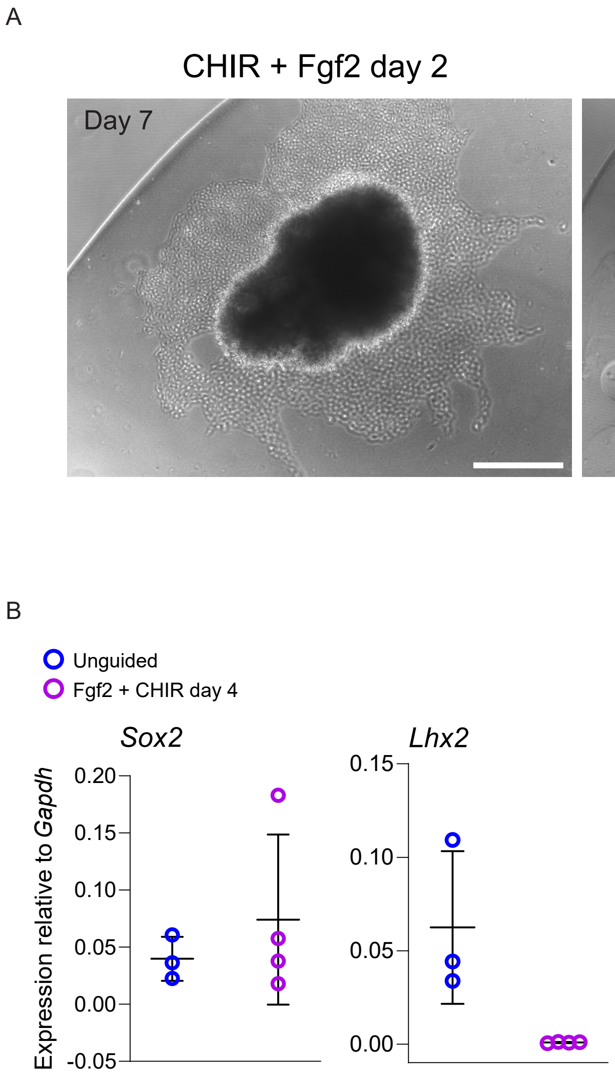
A
CHIR + Fgf2 day 2
CHIR + Fgf2 day 3
Day 7
B
Unguided
Fgf2 + CHIR day 4
Sox2
Lhx2
Expression relative to Gapdh
0.20
0.15
0.10
0.05
0.00
-0.05
0.15
0.10
0.05
0.00

### Supplementary table 1

| | | | |
|---|---|---|---|
| *GBX2* qPCR Fw | GTTCCCGCCGTCGCTGATGAT | Behesti et al., 2021 | human |
| *GBX2* qPCR Rw | GCCGGTGTAGACGAAATGGCCG | Behesti et al., 2021 | human |
| *PTF1A* qPCR Fw | TGAGTTGTTTTTCATCAGTCCA | Watson et al., 2018 | human |
| *PTF1A* qPCR Rw | CAGGCCCAGAAGGTCATC | Watson et al., 2018 | human |
| *KIRREL2* qPCR Fw | GGGGCTAGTTCAGTGGACTAA | Watson et al., 2018 | human |
| *KIRREL2* qPCR Rw | CACGGGCCTAATGTGGAGG | Watson et al., 2018 | human |
| *ATOH1* qPCR Fw | GCGCAAAAGAATTTGTCTCC | Behesti et al., 2021 | human |
| *ATOH1* qPCR Rw | GCGAAGTTTTGCTGTTTTCC | Behesti et al., 2021 | human |
| *GAPDH* qPCR Fw | CACCGTCAAGGCTGAGAACG | Rayon et al., 2020 | human |
| *GAPDH* qPCR Rw | GCCCCACTTGATTTTGGAGG | Rayon et al., 2020 | human |
| *SOX2* qPCR Fw | TGCTGCCTCTTTAAGACTAGGAC | Rayon et al., 2020 | human |
| *SOX2* qPCR Rw | CCTGGGGCTCAAACTTCTCT | Rayon et al., 2020 | human |
| *NANOG* qPCR Fw | GCAACCTGAAGACGTGTGAA | Wamaitha et al., 2015 | human |
| *NANOG* qPCR Rw | CTCGCTGATTAGGCTCCAAC | Wamaitha et al., 2015 | human |
| h*OCT4_POU5f1* qPCR fw | TATGGGAGCCCTCACTTCAC | Wamaitha et al., 2015 | human |

| | | | |
|---|---|---|---|
| h*OCT4_POU5f1* qPCR rw | CAAAAACCCTGGCACAAACT | Wamaitha et al., 2015 | human |
| *LHX2* qPCR Fw | GGGCGACCACTTCGGCATGAA | Ideno et al., 2022 | human |
| *LHX2* qPCR Rw | CGTCGGCATGGTTGAAGTGTGC | Ideno et al., 2022 | human |
| *OTX2* qPCR Fw | ACAAGTGGCCAATTCACTCC | Kirkeby et al., 2017 | human |
| *OTX2* qPCR Rw | GAGGTGGACAAGGGATCTGA | Kirkeby et al., 2017 | human |
| *EN1* qPCR Fw | GGACAATGACGTTGAAACGCAGCA | This study | human |
| *EN1* qPCR Rw | AAGGTCGTAAGCGGTTTGGCTAGA | This study | human |
| *EN2* qPCR Fw | GGCGTGGGTCTACTGTACG | This study | human |
| *EN2* qPCR Rw | TACCTGTTGGTCTGGAACTCG | This study | human |
| *HOXA2* qPCR Fw | CACAAAGAATCCCTGGAAATCG | Wakamatsu et al., 2010 | human |
| *HOXA2* qPCR Rw | AAATGAAATTCTTTTTCCAGCTCTAGA | This study | human |
| *HOXB2* qPCR Fw | GGCCTCTCCCCTAGCCTACA | Dad Abu-Bonsrah et al., 2018 | human |
| *HOXB2* qPCR Rw | GGTGAAAAAATCCAGCTCTTCCT | Dad Abu-Bonsrah et al., 2018 | human |
| *TBR2* qPCR Fw | TAACATGCAGGGCAACAAAATGTATG | This study | human |
| *TBR2* qPCR Rw | GGATTGTAAGACTATCATCTGGGTGTTG | This study | human |
| *PAX2* qPCR Fw | CGTGATGAAGATGTGTCTG | This study | human |
| *PAX2* qPCR Rw | CTGGAAGACGTCAGGGTAGG | This study | human |

| | | | |
|---|---|---|---|
| *SKOR2* qPCR Fw | GTTACCAGTGTCCAAGGCAGAC | This study | human |
| *SKOR2* qPCR Rw | CTGGAAGGCGCTCGACG | This study | human |
| *Gapdh* qPCR Fw | TGACCACAGTCCATGCCATC | Fairchild et al., 2018 | mouse |
| *Gapdh* qPCR Rw | GACGGACACATTGGGGGTAG | Fairchild et al., 2018 | mouse |
| *Skor2 (Corl2)* qPCR Fw | CCCTGTCCACCATCCATC | Muguruma et al., 2010 | mouse |
| *Skor2(Corl2)* qPCR rw | GGTTGTTTTCCTTAGGTTTCTGAT | Muguruma et al., 2010 | mouse |
| *Pax2* qPCR Fw | AAGCCCGGAGTGATTGGTG | Muguruma et al., 2010 | mouse |
| *Pax2* qPCR Rw | CAGGCGAACATAGTCGGGTT | Muguruma et al., 2010 | mouse |
| *Sox2* qPCR Fw | GCGGAGTGGAAACTTTTGTCC | Li et al., 2018 | mouse |
| *Sox2* qPCR Rw | GGGAAGCGTGTACTTATCCTTCT | Li et al., 2018 | mouse |
| *En1* qPCR Fw | GTGGTCAAGACTGACTCACAGC | Aldea et al., 2023 | mouse |
| *En1* qPCR Rw | GCTTGTCTTCCTTCTCGTTCTT | Aldea et al., 2023 | mouse |
| *En2* qPCR Fw | ATGGGACATTGGACACTTCTTC | Soltani et al., 2017 | mouse |
| *En2* qPCR Rw | CCCACAGACCAAATAGGAGCTA | Soltani et al., 2017 | mouse |
| *Gbx2* qPCR Fw | GCAAGGGAAAGACGAGTCAAA | Muguruma et al., 2010 | mouse |
| *Gbx2* qPCR Rw | GGCAAATTGTCATCTGAGCTGTA | Muguruma et al., 2010 | mouse |
| *Lhx2* qPCR Fw | TCGGGACTTGGTTTATCACCT | Xiaodong et al., 2022 | mouse |

| | | | |
|---|---|---|---|
| *Lhx2* qPCR Rw | TGCAAGCGGCAATAGACCAG | Xiaodong et al., 2022 | mouse |
